# High-speed adaptive optics line-scan OCT for cellular-resolution optoretinography

**DOI:** 10.1101/2020.05.20.105478

**Authors:** Vimal Prabhu Pandiyan, Xiaoyun Jiang, Aiden Maloney Bertelli, James A. Kuchenbecker, Utkarsh Sharma, Ramkumar Sabesan

## Abstract

Optoretinography – the non-invasive, optical imaging of light-induced functional activity in the retina – stands to provide a critical biomarker for testing the safety and efficacy of new therapies as well as their rapid translation to the clinic. Optical phase change in response to light, as readily accessible in phase-resolved OCT, offers a path towards all-optical imaging of retinal function. However, typical human eye motion adversely affects phase stability and precludes the recording of fast light-induced retinal events. Here we introduce a high-speed line-scan spectral domain OCT with adaptive optics (AO), aimed at volumetric imaging and phase-resolved acquisition of retinal responses to light. By virtue of parallel acquisition of an entire retinal cross-section (B-scan) in a single high-speed camera frame, depth-resolved tomograms at speeds up to 16 kHz were achieved with high sensitivity and phase stability. To optimize spectral and spatial resolution, an anamorphic detection paradigm was introduced enabling improved light collection efficiency and signal roll-off compared to traditional methods. The benefits in speed, resolution and sensitivity were exemplified in imaging nanometer-millisecond scale light-induced optical path length changes in cone photoreceptor outer segments. With 660 nm stimuli, individual cone responses readily segregated into three clusters, corresponding to long, middle and short-wavelength cones. Recording such *optoretinograms* on spatial scales ranging from individual cones, to 100 μm-wide retinal patches offers a robust and sensitive biomarker for cone function in health and disease. Furthermore, incorporating this capability into an easy-to-use and ubiquitous diagnostic platform of OCT enables its widespread application to patient care and drug development.

## Introduction

Optoretinography, defined generally as the non-invasive, optical imaging of light-induced functional activity in the retina, has the potential to serve as an effective biomarker for retinal health and provide new insight into basic visual processes. Despite its accessibility, the retina presents unique challenges for assessing its physiology in vivo in humans. The retinal reflectivity is low, and the numerical aperture of the eye’s optics further sets a limit on light collection efficiency and lateral resolution of imaging. The challenges are severely compounded due to fixational eye movements that occur at far greater spatial scales compared to the size of retinal neurons. The lack of a non-invasive and sensitive paradigm for the assessment of retinal function poses one of the most fundamental hurdles for developing new therapies and testing their safety & efficacy in humans. Electroretinography (ERG) is the current gold standard, despite the invasiveness of corneal electrodes and severe limits on spatial resolution and signal specificity of the elicited electrical activity^1,2^. Established electrophysiological tools such as patch clamp ^3^ and microelectrode arrays^4^ are not yet feasible for study of retinal physiology in humans. Calcium and voltage-sensitive fluorescent dyes, while extremely powerful as tools for probing cellular physiology in rodent^5^ and non-human primate retina^6^, face regulatory barriers due to potential toxicity. Further, the bright visible illumination required for fluorescence excitation inadvertently stimulates photoreceptors.

Given how optical coherence tomography (OCT), scanning laser ophthalmoscopes (SLO) and fundus cameras are ubiquitous in ophthalmology, incorporating retinal function assessment within these existing imaging platforms would enable their widespread accessibility. Recently, advanced techniques incorporated into OCT, SLO and fundus cameras have been used to visualize spontaneous and light-induced retinal function in vivo in humans^7–14^. Changes in light scattering or intensity^7,12,14–16^, absorption^17–19^, polarization^20^ and optical path length (OPL)^10,21–23^ have been used as all-optical surrogates for function. In addition, variants of motion contrast incorporated into an SLO and OCT serve as an effective biomarker for visualizing blood flow and perfusion^24–27^

Light-induced functional activity in cone photoreceptors has been most accessible, given their higher relative reflectance and amenability for high-resolution *en face* imaging with adaptive optics (AO). Further, the extensive characterization of cone photopigments using genetics, densitometry, psychophysics and electrophysiology^3,19,28,29^ has allowed relating these *intrinsic optical signals* to the underlying mechanisms. For instance, Cooper et al.^12^ recently established that changes in near-infrared image intensity following visible stimuli, such as those observed in AO-SLO^7^ and AO fundus camera^8^ are mediated by the photoisomerization of pigment chromophores. The changes in intensity are presumed to be caused by self-interference of multiple reflections arising from the cone outer segment.

Direct measurements of depth-resolved optical phase changes in response to light are possible in OCT and have provided a sensitive probe for interferometric assessment of light-induced function in the retina. As examples, full field swept source OCT^21^, and point scan swept source^22^, and point scan spectral domain OCT^10,23^ have recently been used for phase-sensitive imaging of light-induced OPL change in the cone outer segments. Each modality must undergo an optimization within the triad of speed, sensitivity and resolution in order to maximize performance. Full-field OCT loses confocal gating across both lateral dimensions, which degrades the x-y spatial resolution and contrast. On the other hand, phase-stable acquisition at high volume rates (hundreds of Hz) with no moving parts are possible that allows computational aberration correction^30,31^. The speed in full-field swept source OCT is ultimately limited by the tunable laser illumination source that must be swept in wavelength at rates that accommodate commercially available aerial sensor frame rates. Swept source and spectral domain point-scan OCT acquire volumes by maintaining confocal gating in both x and y-dimensions that provides excellent resolution and contrast. The speed however is limited by the frequency of the swept laser source, the scanners and the spectrometer, to tens of Hz volume rates. To reach a favorable trade-off in speed and resolution between point scan and full-field OCT, line-scan spectral-domain OCT^32–34^ and line-scan swept-source OCT^35^ have been demonstrated. The line-scan geometry maintains confocal gating along one dimension, while achieving high speeds and phase stability due to the parallel acquisition of multiple A-lines simultaneously. Specifically, in line-scan spectral domain operation, one B-scan is captured per 2D camera frame. Ginner et, al.^34^ demonstrated its application for computational aberration correction and Zhang et al.^33^ incorporated AO, to improve resolution in retinal cross-sections, with maximum B-scan rates of 2500 Hz and 500 Hz respectively. In addition, Ginner et al. quantified the median axial velocity across 36 subjects and concluded that a B-scan rate greater than 3200 Hz was necessary to avoid greater than ±*π* radian phase change for interferometric applications.

Here we present the first high-speed AO line-scan OCT for volumetric imaging and phase-resolved acquisition of light induced retinal activity. We detail the design, development and characterization of the multimodal imaging system. The feasibility of imaging light induced OPL changes, both with and without AO in cone outer segments at high temporal resolution is demonstrated. Measurement of such light induced optical changes in the retina are referred to as an optoretinogram (ORG) in analogy to the classical ERG. This terminology was first proposed by Mulligan et.al.^36,37^ and adopted recently by our group and others to refer to stimulus induced optical imaging of retinal responses using OCT phase and intensity signatures^38–40^. Looking ahead, we suggest that the ORG terminology be used independent of the platform (OCT, SLO or fundus camera), type of optical change (intensity or phase) or spatial scale (individual cells to collection of neurons) to refer to light-induced changes in the retina in general.

## Results

### System layout and specifications

A free space, multimodal, line-scan retinal imager consisting of a line-scan spectral domain OCT and line-scan ophthalmoscope (LSO) was designed and constructed. Three illumination and detection channels were incorporated. A superluminescent diode (SLD) (M-S-840-B-I-20, Superlum, Ireland, λ_o_ = 840 nm, Δλ = 50 nm, axial resolution = 7.6 μm in air) was used as illumination for OCT and for LSO, a 980±10 nm SLD (IPSDD0906, Inphenix, USA) served as illumination for wavefront sensing, and a 660±10 nm LED illuminated a ~37.5 deg^2^ retinal area in Maxwellian view for visual stimulation. Three detectors, one each for OCT, LSO and a Shack-Hartmann wavefront sensor captured the backscattered light from the retina.

The basic principle behind the operation of an OCT in line-scan spectral domain configuration involves illuminating the sample with a line-field, achieved here with a cylindrical lens. In detection, the backscattered linear spatial profile from the retina is made to interfere with a reference beam. The interference is then diffracted into its spectral constituents with a linear diffraction grating, yielding a two dimensional image on an aerial sensor, corresponding to the spatial and spectral components along the two axes. Traditional OCT processing provides a cross-sectional retinal image or B-scan in one aerial camera frame, the rate of which is limited by the maximum camera frame rate. Here, a high-speed CMOS camera was used that allowed recording a maximum B-scan rate of 16 kHz, for a region-of-interest of 768 (spectrum dimension, λ) ×512 (line dimension, *x*) pixels. The spectral resolution corresponded to an axial range of 2.6 mm. For volumetric imaging, the line-field was scanned using a 1-dimensional scanner. Both non-AO and AO modes of imaging were used. A deformable mirror (DM) in the sample arm corrected the aberrations of the eye. An adjustable pupil iris in the sample arm was set to 4 mm to reduce the effect of eye’s aberration in non-AO mode of imaging, where defocus was optimized. For AO imaging, the adjustable iris was opened fully, such that the DM aperture formed the limiting pupil, corresponding to 6.75 mm at the eye. The detection path consisted of two additional features. First, an anamorphic configuration provided asymmetric lateral magnification that enabled the optimization of spectral and spatial resolution, while simultaneously improving light collection efficiency. Second, the zeroth-order diffraction beam from the grating, which is typically discarded in spectral-domain OCT systems, was used to build an LSO by placing a line-scan camera optically conjugate to the OCT camera. The LSO enface real-time image stream allowed precise adjustment of the image focus. The optical layout of the system is described in detail in Materials and Methods.

### Anamorphic detection

In traditional point-scan spectral domain OCT, the spectrometer – grating, focusing lens and line-scan camera – are chosen in order to optimize spectral resolution or axial range. Of these parameters, the line-scan camera pixel size and grating choices are restricted to those available from commercial suppliers. Therefore, the focusing lens focal length is optimally selected to obtain the desired spectral resolution for a given source spectral bandwidth.

In addition to spectral resolution, the detection optics in line-scan spectral domain OCT need to achieve optimal spatial sampling along the line/spatial dimension in order to resolve fine retinal structures such as photoreceptors. The spatial resolution depends on the overall system magnification and thus clearly differs in its governing parameters compared to those limiting spectral resolution. To achieve an optimal trade-off in spectral and spatial resolution necessitates independent control over the system magnification in both lateral dimensions, such that the square pixel sampling of typical aerial cameras is sufficient. Conventional spherical lenses do not offer sufficient degrees of freedom to optimize these factors independently. An anamorphic configuration consisting of two cylindrical lenses offers a solution to optimize magnification asymmetrically. In an LSO, Lu et al.^41^ demonstrated that such an anamorphic telescope allowed adequate spatial sampling and improved signal collection efficiency. For line-scan spectral domain OCT, we posited that such a configuration will reduce cross-talk between adjacent pixels in the spectral dimensions, significantly improving signal roll-off as a result.

Figure 1 shows the anamorphic detection layout and performance. In Figure 1a, the image plane I_circ_ prior to spherical achromat L_A_ contains spherically symmetric point-spread-functions (PSFs) at each field point. Once collimated, the beam profile at the system’s exit pupil plane (P_circ_) is also spherically symmetric. Next, a pair of positive cylindrical lens achromats CLx (EFL = 250 mm) and CLy (EFL = 100 mm) were placed, with their power axes oriented orthogonal to each other. The distance between the two lenses was set to the difference of their focal lengths. At the imaging plane, I_anamorphic_, an elliptical PSF is obtained, whose major and minor axes sizes correspond to the ratio of the cylindrical lens focal lengths, equal to 2.5. Once the light from this imaging plane is collimated with a spherical lens (L_B_), the pupil plane P_anamorphic_ containing the grating is similarly elliptical in shape, with the same ratio of its major and minor axes. Effectively, this configuration provides 2.5× higher sampling in the horizontal (spectral) compared to the vertical (spatial) dimension.

**Figure 1:**
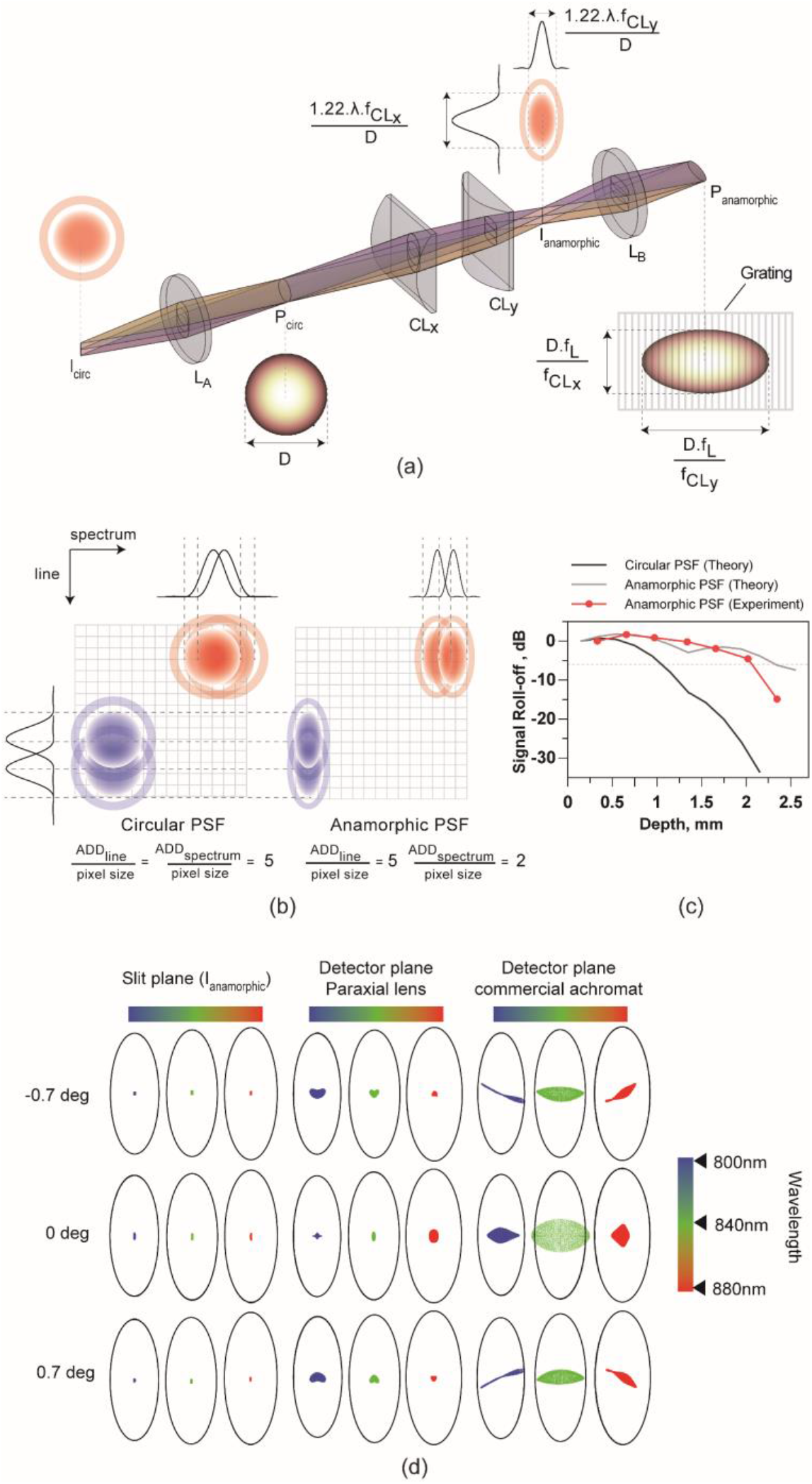
Anamorphic detection to optimize spatial and spectral resolution (a) Projection view (trimetric) of the anamorphic detection scheme for line-scan ophthalmoscopy. Note that the figure shows singlet lenses for simplicity, however all lenses were achromatic doublets. The pupil plane (P_circ_ and P_anamorphic_) and imaging plane dimensions (I_circ_ and I_anamorphic_) scale along their axes depending on the cylindrical lens (CLx and CLy) focal lengths. (b) The circular and anamorphic PSFs and their 1-dimensional cross-sections are shown along the line and spectral dimensions, overlayed on the aerial detector pixels, illustrated to scale. The number of camera pixels contained within one airy disk diameter (ADD) along the line and spectrum dimensions is indicated for both circular and anamorphic PSFs. Along the line dimension, the spatial sampling is identical between both as evident in the 1-dimensional cross-section, however, the anamorphic PSF offers increased light collection per pixel. Along the spectrum dimension, the resolution is increased by 2.5× in the anamorphic PSF in comparison to the circular PSF, as evidenced by the greater separation in the 1-dimensional cross-section. (c) Signal roll-off versus depth is shown to demonstrate the effect of the spherical and anamorphic PSFs. Also shown is the measured signal roll-off for our anamorphic detection spectrometer that shows good correspondence with theoretical computation and demonstrates the substantial improvement over circular PSFs. (d) The theoretical imaging performance of anamorphic detection represented as spot diagrams for different field points and wavelengths along with the diffraction limited airy ellipses for reference. The spot diagrams at three imaging planes are shown, corresponding to (i) the imaging plane after the cylindrical lenses (I_anamorphic_) prior to diffraction via the grating (ii) detector plane, with an ideal paraxial lens as the spectrometer focusing element to demonstrate the effect of the cylindrical lens telescope plus diffraction, on imaging performance and (iii) detector plane with the used commercial achromatic doublet as the focusing optic to demonstrate the effect of all optics on imaging, as realized practically in the system.

In principle, the focal lengths of CLx and CLy can be chosen according to the camera, grating and source bandwidth in consideration; here chosen as such to provide a Nyquist limited spatial resolution of 2.4 μm × 2.4 μm and a theoretical axial range of 2.6 mm. In Figure 1b, consider the difference between a spherical and an anamorphic PSF in optimizing resolution, signal collection and crosstalk in both dimensions. For the line/spatial dimension oriented along the major axis of the elliptical PSF, the anamorphic PSF allowed increased signal collection per pixel, in addition to adequate spatial sampling. A significant improvement is expected for the spectrum dimension oriented along the minor axis of the elliptical PSF, because of the reduced cross-talk between adjacent pixels. The most appreciable advantage of the reduced spectral cross-talk is expected in the improved OCT signal roll-off performance (Figure 1c), provided the PSF is diffraction-limited for all field and spectrum points (Figure 1d).

The imaging performance of the anamorphic detection was modeled with respect to the spatial and spectral dimensions. Signal roll-off under conditions of a spherical and an anamorphic PSF were theoretically computed. For the spherical PSF, the diffraction-limited airy disk diameter (ADD) encompassed ~5 camera pixels along both dimensions. For the anamorphic PSF, the ADD was reduced to 2 pixels in the spectral dimensions, while the ADD in the spatial dimension was same. These PSF dimensions were computed based on the spectrometer lens focal length, camera pixel size and the beam size at the pupil plane containing the grating. First, the interference modulation in wavelength space was created as a function of OPL or axial depth and superimposed with the measured source spectrum. The interference modulation was convolved with the circular and elliptical PSFs to reflect the effect of diffraction due to the finite size of the PSFs itself. In principle, the PSF dimensions at each spectral position will essentially be weighted by the wavelength, but this wavelength-dependent variation for the short 50 nm bandwidth source used here was not taken into account for this calculation. Instead, all PSFs were computed for a center wavelength equal to 840 nm.

Once convolved, the interferograms were subjected to traditional OCT reconstruction – k-space remapping, Fourier transform and mirror image removal. The peak signal at each axial depth is plotted in decibels (dB) in Figure 1c for both the anamorphic and spherical PSFs. The improvement in signal roll-off is evident for the anamorphic PSF with this theoretical treatment. Figure 1c also shows how the measured signal roll-off agrees well with theory for the anamorphic detection. The 6 dB signal roll-off occurred experimentally at 2.1 mm. At 1.5 mm and 2.1 mm axial depths, the anamorphic PSF outperforms the circular PSF by 10.2 dB and 29.8 dB, respectively.

Figure 1d shows the spatial imaging performance across different discrete fields and wavelength points of the system modeled in ZEMAX optical design software. Note that the ellipses in Figure 1d spot diagrams denote the diffraction-limited “Airy ellipses” (as opposed to Airy disks), because the diameters of their major and minor axes are determined by the powers of cylindrical lenses. The ZEMAX model started from the pupil plane P_circ_, with the specified imaging field points and wavelengths. It contained the rest of the optical elements in the detection path (CLx, CLy and L_B_), plus a transmissive 1200 lp/mm grating and a focusing lens (EFL = 200) prior to the camera image plane. The distances between CLx and CLy, and between the focusing lens and the image plane were optimized in the model to provide minimum spot sizes at I_anamorphic_ and camera sensor, respectively. Three cases are shown to separate the contributions due to the cylindrical lens telescope, diffraction due to the grating and the achromatic doublet spectrometer focus lens. The first column corresponds to the spot diagrams at the slit image plane I_anamorphic_ where performance is diffraction-limited. The spot diagrams after diffraction through the grating and focused on the camera by an ideal paraxial lens is shown in the middle column, demonstrating diffraction-limited performance also. The last column shows the same, but with the off-the-shelf achromatic doublet focus lens used in the system. The field curvature typically observed in spectrometer designs is apparent with the doublet. Despite the use of an off-the-shelf achromat, the spot sizes for all the field and wavelength points, except 0 deg/840 nm, are diffraction-limited. A custom-design achromat or a lower grating pitch leading to a smaller Bragg angle of diffraction ought to improve performance further for such an anamorphic configuration. Overall, anamorphic detection improves spectral resolution, signal roll-off and light collection efficiency, without adversely affecting imaging resolution. Further, it allows customizing camera regions-of-interest for increased acquisition speeds, and the use of commercially-available parts, i.e. transmissive gratings and achromatic focusing optics.

### System characterization

The system performance was characterized with respect to sensitivity, axial resolution, signal roll-off and phase sensitivity.

#### Sensitivity and Signal roll-off

The sensitivity of the system was measured by imaging OCT B-scans with a flat mirror and a neutral density filter placed in the sample arm, at the plane corresponding to eye’s pupil. The axial position of the reference mirror was set to 315 μm in depth. The OCT interferograms were subjected to k-space remapping, Fourier transform and mirror image removal to retrieve the SNR in dB. The system sensitivity was obtained by adjusting the SNR for the neutral density filter attenuation. This process was repeated at different exposure times/frame rates, ranging from 62.5 to 400 μsec (2.5 to 16 kHz). For shot-noise limited performance, a linear increase in sensitivity is predicted with integration time. The collimator and cylindrical lens focus together induce a Gaussian distribution of intensity along the linear illumination. Therefore, the SNR follows the same distribution along the line, the variation in magnitude of which is indicated by the error bars at each frame rate in Figure 2a. Here, for the lowest frame rate of 2.5 kHz, the sensitivity was 89.6±2.2 dB and decreased linearly with log of integration time (Figure 2a). Sensitivity was also measured as a function of depth to retrieve the signal roll-off (Figure 1c).

**Figure 2:**
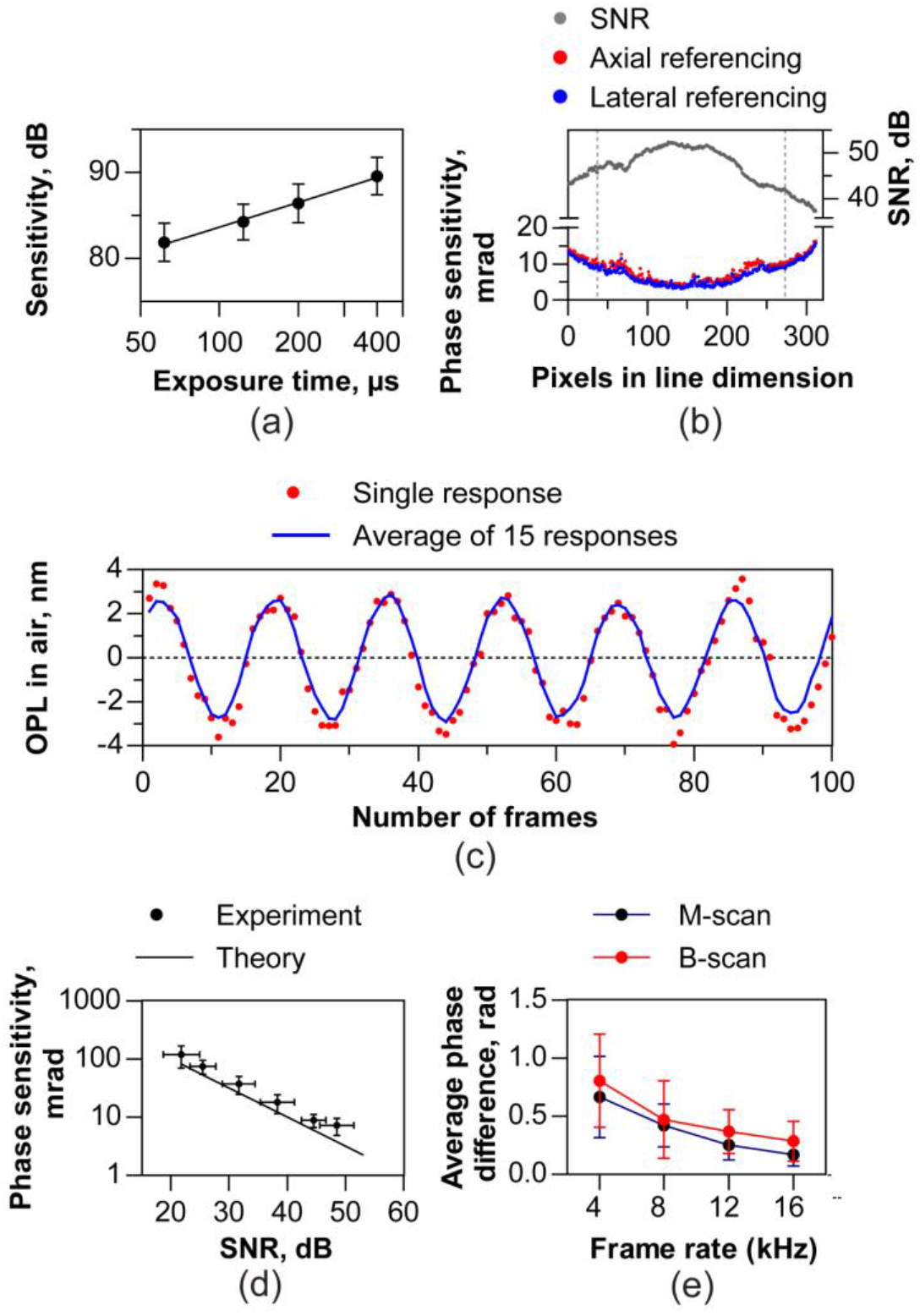
Signal-to-noise and phase sensitivity characterization (a) Sensitivity in dB varies linearly with log of exposure time of the aerial sensor indicating near shot noise limited performance. (b) Measured phase sensitivity (bottom panel) in milliradians (mrad) along the line illumination for a coverslip. The axially referenced (front and back surface of cover slip) and laterally referenced (two points separated on the line) phase sensitivity is plotted and are similar. The phase sensitivity is worse at the edges of the line profile consistent with the lower SNR (top panel) resulting from a Gaussian intensity distribution along the line illumination. (c) Optical phase change represented in terms of optical path length in air resulting from a piezo-transducer held under a sinusoidal voltage signal. The phase was calculated by laterally referencing two points along the line, spaced by 75 pixels. A single response and the mean of 15 responses show good agreement. (d) Measured phase sensitivity versus SNR follows a linear relationship and is close to the theoretical shot-noise limited phase sensitivity, defined as the square-root of SNR^-1^. The data points and error bars in (a) and (d) denote the mean and standard deviation sensitivity, SNR and phase noise for 236 pixels along the line, corresponding to 70% of the spatial extent of the Gaussian illumination (vertical dashed lines in Figure 2b). (e) Axial bulk motion quantified as average phase difference in the two modes of operation – M-scan and multiple B-scans per volume, decreases monotonically with increasing frame rate. The error bars denote the mean variability, calculated as the mean of the standard deviation across the 10 measurements for each speed.

#### Phase sensitivity

Phase sensitivity (*σ*_Δ*φ*_) in OCT is intimately linked to SNR, motion (Δx) of the sample, and the spot size (‘d’)^42,43^ as shown in the equation 1 below.

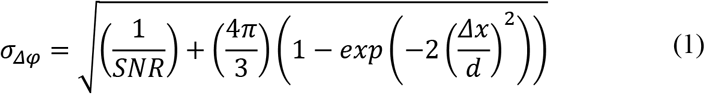

Therefore, it is instructive to quantify phase sensitivity as a function of SNR and sample/eye motion to understand the systematic limits for interferometric applications. First, the phase sensitivity was measured for different SNRs using an artificial model eye with a 50 mm achromat focus lens and a coverslip as the retina. M-scans, i.e. B-scans at the same location versus time were recorded by varying SNR using a variable neutral density filter in the sample arm. A set of thousand M-scans were reconstructed, and the phase difference between the front and back surface of the cover slip was determined for a single point over time. This denoted the axial referencing method for computing phase sensitivity. Note that in line-scan operation, the parallel acquisition of a B-scan allows lateral referencing (also called self-phase referencing ^44^) to remove common mode noise. For lateral referencing, the phase difference between two immediately adjacent pixels on the front surface of the cover glass was measured. This was repeated for each pair of adjacent pixels ranging across the length of the line illumination. A measure of phase sensitivity or stability is given by the standard deviation of the phase differences measured via either axial or lateral referencing. The effect of the galvo scanner jitter was quantified by measuring axially referenced phase sensitivity under three conditions: a) galvo powered off, b) galvo powered on, but actively reset to zero volts and c) a linear galvo scan across 250 locations in a 0.75 deg field of view.

Figure 2b shows the measured phase noise and SNR corresponding to the highest sample power for the ~300 pixels along the line dimension. The Gaussian intensity profile along the line illumination reduced SNR at the edges, leading to an increase in phase noise. At all points along the line, the axial and lateral referencing produce similar phase noise, limited to less than 15 mrad at this sample power. The Gaussian variation in SNR and phase noise across the line field is captured by error bars in Figures 2a and 2d. Figure 2d shows the relation between the SNR and phase sensitivity – a linear decrease in phase sensitivity vs. SNR is observed. Upon setting the dependence of phase sensitivity on motion, the term ‘Δx’, to zero, in equation 1, the shot-noise limited phase sensitivity for a static sample reduces to the reciprocal of the square root of SNR (theoretical curve – solid line – in Figure 2d). The good match between the measured and theoretical phase noise demonstrates that the system phase sensitivity is near the shot-noise limit. For the highest sample power, a mean phase sensitivity of 7.5±2.3 mrad was measured along the line with the galvo turned off. With the galvo actively set to zero volts, phase sensitivity was similar as when it was turned off, equal to 7.2±2.3 mrad. However, scanning a linear 0.75 deg field, the galvo jitter increases phase noise substantially by ~10 times to 76.6±38 mrad. On the other hand, the variation in phase noise across the line illumination was uniform *and* low, less than 15 mrad at the same sample power. This demonstrates a key advantage in phase stability for parallel acquisition in line-scan geometry and how moving parts, i.e. scanners can adversely affect phase noise.

To further test the ability of laterally referenced phase differenced measurements to quantify axial change, we replaced the coverslip with a piezoelectric transducer at the focal plane of the model eye. The piezoelectric transducer deforms in shape according to the drive voltage. The amplitude and frequency of axial physical displacement was controlled with a sinusoidal voltage waveform. We sought to calculate the smallest axial displacement measurable on the surface of the transducer. The phase difference was calculated between two points separated laterally on the surface of the transducer by 75 pixels, converted to OPL change and plotted versus the number of camera frames in Figure 2c. A minimum amplitude of 3 nm was observed in a single response that was repeatable across 15 repeats.

Axial bulk motion adversely affects phase sensitivity and can be quantified by measuring the phase difference between two consecutive B-scans (see Figure 3 for more details). We observed that the average phase difference decreases monotonically with increasing acquisition rate (Figure 2e). The average phase difference shown in the Figure 2e were obtained for 10 sets of measurements at each speed for M-mode and volumetric acquisition. The mean and standard deviation axial bulk motion for volumetric acquisition is expectedly higher, due to galvo jitter, but not statistically significant.

**Figure 3:**
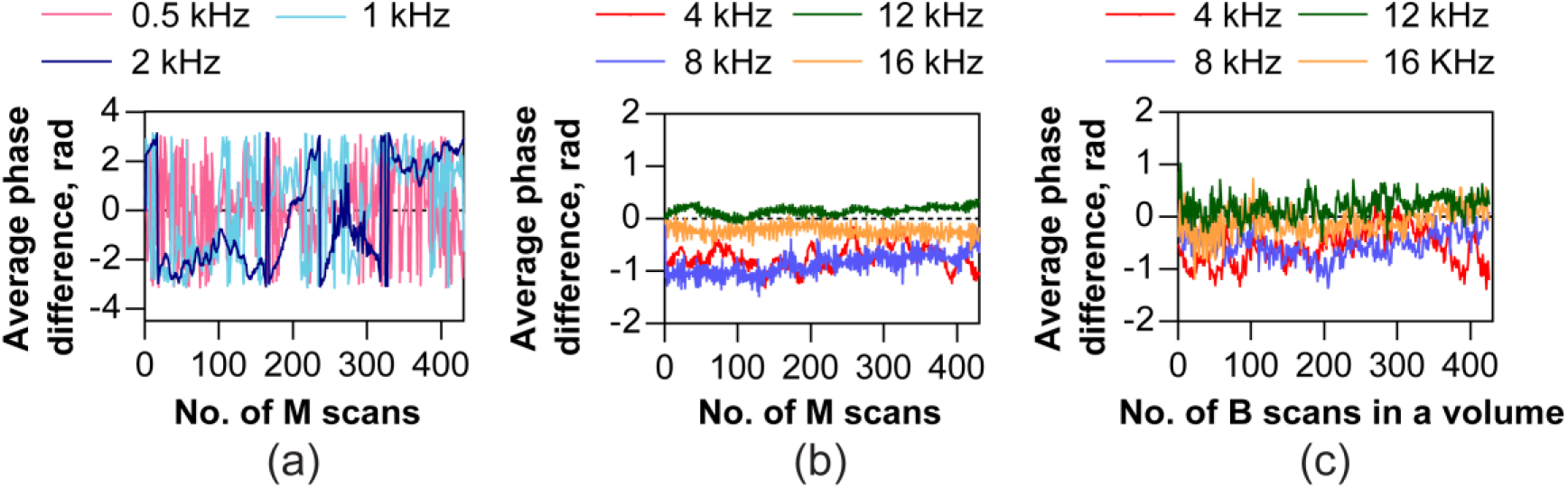
Axial bulk motion for a range of M-scan and B-scan rates. Average phase difference between successive M-scans with (a) 0.5 kHz, 1 kHz and 2 kHz frame rates show severe phase wrapping that prevents its optimal correction and with (b) 4 kHz, 8kHz, 12 kHz and 16 kHz frame rates show no phase wrapping enabling accurate correction of axial bulk motion. (c) The average phase difference between successive B-scans along the scan direction in a volume shows no phase wrapping when measured in the range between 4 kHz – 16 kHz B-scan rates.

##### Effect of eye motion on phase sensitivity at different B-scan rates

To compensate axial bulk motion reliably, the resulting maximum phase change should ideally lie within ±*π* radians. Here, we estimated the average phase change between two consecutive B-scans recorded at different speeds ranging from 0.5 to 16 kHz. M-scans were recorded with the galvo set to 0 volts and volumes were recorded with the galvo scanning, corresponding to an overall field of view equal to 2 × 3.5 deg. Average phase difference due to axial bulk motion was compared in M-mode and volumetric imaging across different camera frame rates. The complex phase in the (n+1)^th^ B-scan was averaged and multiplied with the complex conjugate of the average complex phase in the n^th^ B-scan. For M-mode, ‘n’ signifies the number of B-scan acquisitions at the same spatial location, while for volumetric acquisition, ‘n’ signifies the number of scans. These measurements were made without AO, for a 4 mm pupil with defocus optimized.

In Figure 3a, B-scan rates less than 2 kHz show substantial phase instabilities, leading to phase wrapping between consecutive B-scans. Such phase drifts during a single B-scan acquisition are attributed to eye motion and lead to fringe washout and reduced SNR. Fringe washout is a well-recognized issue in spectral domain OCT operation while acquiring A-scans at low speeds, evident here in line-scan geometry for low B-scan rates. This observation aligns well with Ginner et al.^34^ For B-scan rates in the range between 4 - 16 kHz, we observed that the average phase difference between consecutive B-scans remained below ±*π* radians, both in M-mode (Figure 3b) and volumetric (Figure 3c) acquisition.

### Imaging retinal structure

Figure 4 shows retinal images taken with AO with the LSO and OCT mode of operation. The focus was optimized at the outer retina to reveal cone photoreceptors. Figures 4 a-b & c-f show the volume and cross-sectional images obtained at 4 and 2 deg temporal eccentricity. From the AO-corrected volumes (Figures 4a & b), the substantially higher signal in the outer retina compared to the inner retina is indicative of the small depth of field of imaging, resulting from the larger numerical aperture with AO. The LSO images in Figures 4c and 4e were obtained at a high integration time, and low speed of 30 Hz. This was to account for the limitation in signal, arising by placing the LSO focus optic and detector at the zeroth order of the diffraction grating, accounting for ~10 % of the light in the detection arm. Figures 4d and 4f represent maximum intensity projections obtained at the cone outer segment tips (COST). The OCT volumes were recorded at 5000 B-scans/sec (10 vols/sec – 500 scans per volume). In both LSO and OCT *en face* images, cone photoreceptors were clearly resolved. At eccentricities closer to the fovea, imaging resolution was restricted by Nyquist sampling of detection, optimized as such to favor speed and light collection efficiency per pixel.

**Figure 4:**
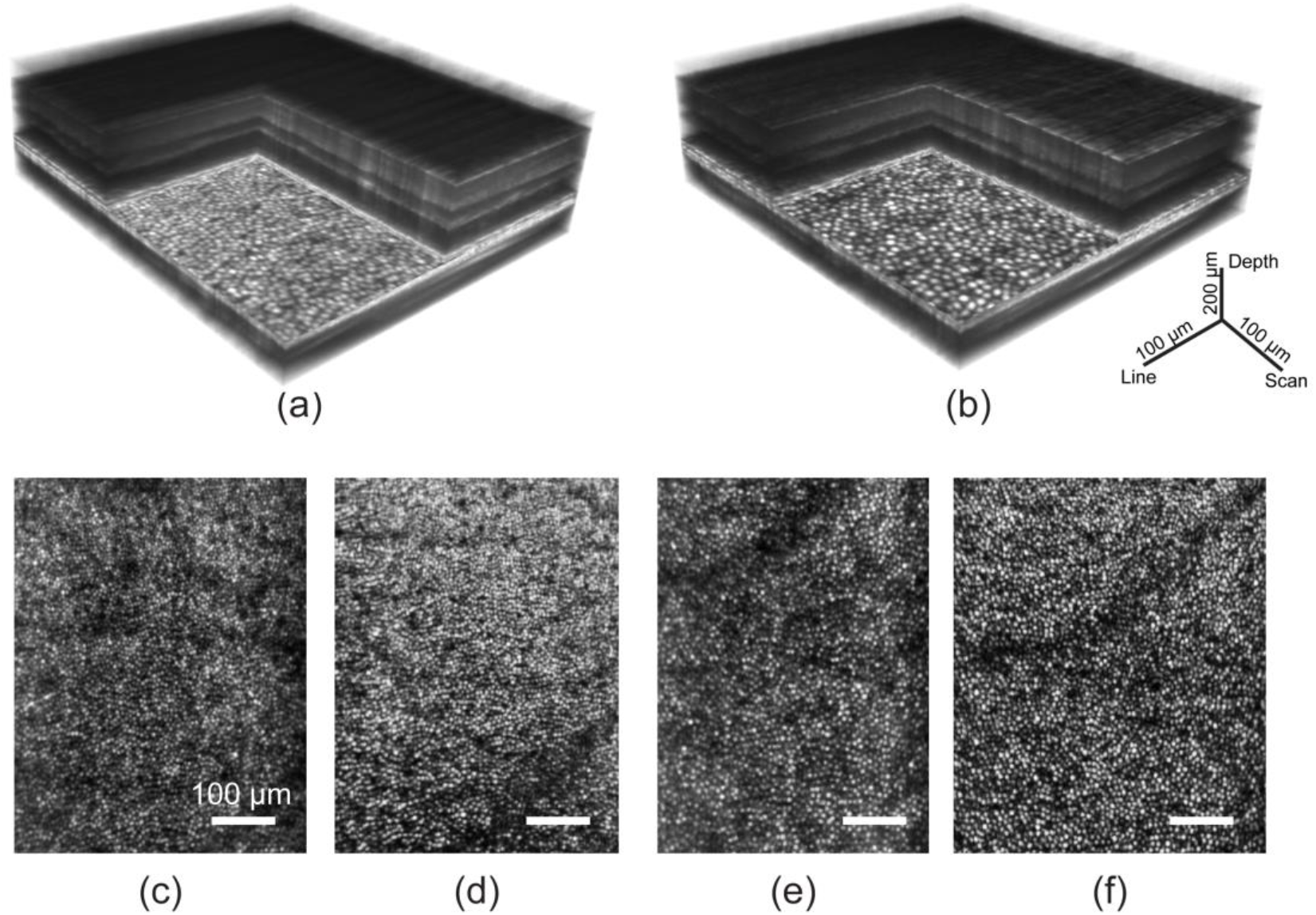
High-resolution imaging of retinal structure with AO line-scan spectral domain OCT and line-scan ophthalmoscope. Three dimensional rendering of AO-OCT volumes acquired at 2 and 4 deg temporal eccentricity are shown in (a) and (b). The corresponding en face images at the COST layer show cone photoreceptors in (d) and (f). LSO en face images are shown in (c) and (e). Scale bars for the volumes are indicated and are equal to 100 μm for the enface images.

### Optoretinography without AO

The OCT volumes were segmented for the layers corresponding to cone inner-outer segment junction (ISOS) and COST. The phase at the ISOS and COST layers were axially referenced (*ϕ*_*COST*_ − *ϕ*_*ISOS*_) with respect to each another to denote the light-induced OPL change in the cone outer segment after 660 nm light stimulus, and plotted versus time to reveal its dynamics. The change in OPL obtained from the phase difference (*ϕ*_*COST*_ − *ϕ*_*ISOS*_) was averaged over 0.27 deg^2^ on the retina. Each curve represents an optoretinogram or ORG, and was obtained without AO for a 4 mm pupil and defocus optimized. Figures 5a and 5b were obtained at volume rates of 324 Hz and 120 Hz respectively to reveal light induced OPL dynamics in cone photoreceptors across different temporal scales.

**Figure 5:**
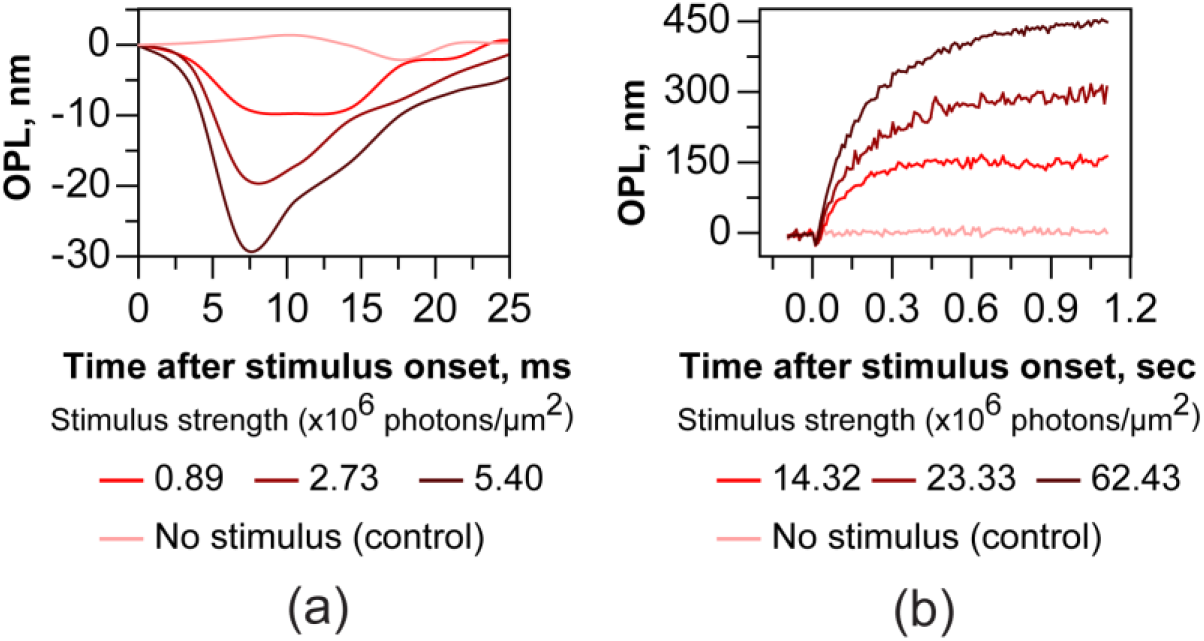
Optoretinograms with 660 nm stimulus acquired without adaptive optics. After stimulus onset, cone outer segments undergo a rapid shrinkage in OPL (a) followed by a slower expansion (b). The response magnitude and rate of change scale with increasing stimulus strength. The phase response was averaged over a retinal area of 0.27 deg^2^. In a), each curve is an average of two separate measurements, while single measurements are shown in b). The early reduction (ORG early response) and later expansion (ORG late response) were measured at 324 Hz and 120 Hz respectively. For the control (no stimulus case), the mean ± standard deviation OPL calculated over ~1.2 seconds was 2.2 ± 4.9 nm for the single measurement.

In Figure 5a, a rapid reduction in OPL immediately after stimulus onset is observed. The maximum amplitude of shrinkage in OPL was 9 nm, 19 nm and 29 nm for the three stimulus intensity levels. Only two separate measurements were averaged for the traces in Figure 5a to reduce phase noise for these minute OPL magnitudes recorded at high speed. In Figure 5b, while the decrease is not readily apparent at the lower speed, an increase in OPL is observed after stimulus onset, saturating at 152 nm, 294 nm and 444 nm, ~500 ms post-stimulus. While the fidelity of these curves will certainly improve with averaging repeat measures, the curves in Figure 5b represent single unaveraged recordings in order to establish the limits of phase stability in the line-scan paradigm. For the control (no stimulus case), the mean ± standard deviation OPL calculated over ~1.2 seconds was 2.2 ± 4.9 nm for the single measurement (Figure 5b). In general, the amplitude of the initial reduction and later expansion of OPL scales with the stimulus energy, tested in the range between 14.3×10^6^ to 62.4×10^6^ photons/μm^2^ for the late expansion and 0.9×10^6^ to 5.4×10^6^ photons/μm^2^ for the early reduction.

The scaling of the ORG response – both the reduction and increase in OPL – with stimulus energy has been demonstrated before using point-scan spectral domain and swept source OCT. The late response has been attributed to be the influx of water to maintain osmotic balance during the phototransduction cascade ^15^. Based on the latency and the magnitude of the stimulus strength that characterize the rapid shrinkage, we earlier concluded that the reduction in OPL is the optical manifestation of the early receptor potential observed previously in vitro, attributed to charge movement across the outer segment disc membranes during photoisomerization^39,45–47^.

### Optoretinography with AO

Similar experiments as Figure 5 were repeated with the increased resolution offered by AO that allowed discerning the ORG in individual cones, in response to the 660 nm stimulus. Figure 6a shows a cone photoreceptor image at 4.75 deg eccentricity obtained with AO-OCT. The mean ± std of the ORG in individual cones (n = 450 cones) is plotted in Figure 6c. The ORG maximum amplitude and time course is variable across cones. The histogram of the response amplitude averaged between 0.5 to 1.2 seconds was subjected to Gaussian mixture model clustering analysis (Figure 6d). Three clusters emerged from this analysis, where each cone belonged to a sub-group with high probability (mean ± std: 0.99±0.05 across all cones) and low uncertainty (< 0.18%). For a cone within each sub-group, the probability is defined as the ratio of its component Gaussian to the sum of all Gaussians, while the uncertainty is defined as the area of overlap between the component Gaussians. Based on the intersection of the component Gaussians, these clusters were labelled as those belonging to S, M and L-cone photoreceptors as shown in Figure 6b, ordered by their decreasing sensitivity to 660 nm wavelength. In Figures 6b-d, the color of individual cones and their corresponding ORGs are labelled as ‘red’, ‘green’ or ‘blue’ to denote the three cone types, on the basis of their spectral identity obtained from the clustering analysis. The maximum amplitude (mean ± std) of L and M cones was 228±40 nm, 90±26 nm, corresponding to a photopigment bleach of 21.8% and 3.8 % respectively. At 660 nm wavelength, S-cone photopigments are bleached to a negligible degree (<0.001%) resulting in an insignificant OPL change of 12±22 nm.

**Figure 6:**
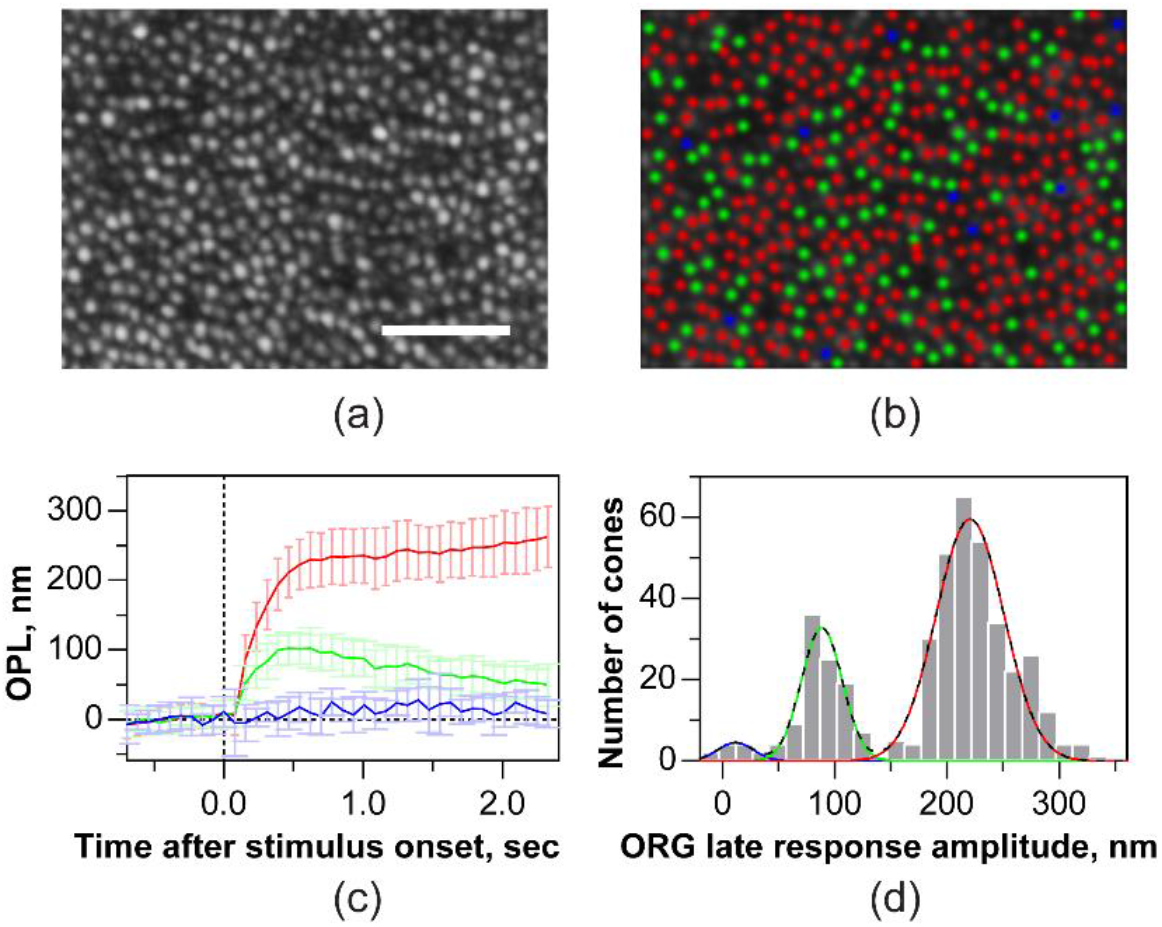
Single cone optoretinograms (ORGs) with 660 nm stimulus acquired with AO. (a) Cone photoreceptor image at 4.75 deg temporal eccentricity obtained at the COST layer from a set of registered AO-OCT volumes. (b) Long-, Middle- and Short- (LMS) cones marked as ‘red’, ‘green’ and ‘blue’, classified based on Gaussian mixture model clustering of the ORG magnitude shown in (d). (c) the mean and standard deviation of the ORG in individual L-, M- and S-cones. Solid lines and error bars denote the mean and ± 1 std across the population of cones. (d) A histogram showing the distribution of ORG magnitude. The black dashed line denotes the sum of three Gaussians obtained from the clustering analysis. The individual component Gaussians are shaded red, green, blue, denoting the LMS-cones respectively. Volume rate was 15 Hz. Scale bar: 50 μm.

## Discussion and Conclusions

The line-scan spectral domain operation with AO offers an excellent opportunity to optimize the triad of speed, sensitivity and resolution and would benefit applications of OCT ranging from angiography to clinical retinal structure-function imaging. The B-scan rates up to a maximum of 16 kHz, to the best of our knowledge, represents the highest speeds for two-dimensional cross-sectional OCT imaging of the retina. This speed is limited by the maximum camera frame rate. The camera cost is higher than conventional CCD and CMOS sensors, though at ~$40,000, our sensor is relatively less expensive than some science-grade CCDs and still faster burst-mode cameras, that can cost up to ~$100,000. The camera provided fast phase stable acquisition for multiple effective A-lines in parallel and is thus essential for resolving the rapid light-induced retinal responses, that may otherwise by severely confounded by eye motion. Specifically, volume acquisition rates as high as 324 Hz was required to resolve responses with short latency and fast dynamics such as the early axial contraction of outer segments. Alternatively, analyzing the phase at the resolution of individual B-scans, either acquired in succession within one volume, or in separate volumes, provides increased temporal resolution at the cost of complete loss in spatial information^23,39^. We summarized the relationship between acquisition speed and axial bulk motion for in vivo imaging across a large temporal range extending measurements of Ginner et al. up to 16 kHz B-scan rates. Given the impact of axial bulk motion on fringe contrast in spectral domain OCT and its application to interferometric imaging, this quantification will allow future optimizing of speed and sensitivity for line-scan operation.

An added benefit of parallel phase-stable acquisition is the ability to perform lateral or self-phase referencing. The optical phase at any given single point in time and space by itself is essentially noisy due to both light-tissue interaction and external factors such as air currents, galvo jitter and eye motion. Thus, overcoming common mode noise is essential to reveal robust OPL changes. This is performed by a combination of referencing the phase in space along the axial direction – for instance COST vs. ISOS; and/or referencing phase in time – with stimulus vs. without stimulus. We show a demonstration of lateral phase referencing using a piezotransducer possible due to parallel cross-sectional imaging. While the minimum phase change detectable is still restricted ultimately by SNR, lateral phase referencing enables measuring OPL change across two laterally separated points in space, two neighboring cones or blood vessels for example.

The parallel phase-stable acquisition comes at the cost of loss of confocality across one lateral dimension which affects the imaging contrast. The anamorphic detection configuration that we introduce for line-field spectral domain OCT, while optimizing spatial and spectral resolution independently, also simultaneously increases light collection efficiency per pixel and as result, the contrast. The benefits of the anamorphic configuration are evident in the improved signal roll-off and the Nyquist-limited imaging of cone photoreceptors. In addition, this is achieved by the use of commercially available parts allowing efficient future implementation. We demonstrate the feasibility of recording light-induced responses across small retinal areas without AO (defocus optimized) and in individual cones with AO. The former is encouraging as an avenue to perform optoretinography over larger fields of view and smaller undilated pupils in a clinical population and underlines the feasibility of such recordings without the use of additional hardware for AO. With AO, we revealed that the optoretinograms can be assigned to individual cones that enables their segregation into L, M and S-cones based on the magnitude of their response to 660 nm stimulus. In comparison to AO retinal densitometry^18,19^, this technique of classifying the cone spectral types provides not only exceedingly higher precision but also considerably shorter imaging durations, as characterized by Zhang et al. using a point-scan spectral domain AO-OCT^23^.

Some shortcomings remain to be addressed in future work. First, line-scan geometry may cause multiple scattering artifacts for imaging structures near and distal to the pigment epithelium. We have not measured here the extent to which multiple scattering affects the fidelity of imaging and remains an active area of investigation. Second, the cylindrical lens used to generate a line illumination maintains the Gaussian intensity distribution, resulting from fiber collimation, across the imaging field. Phase sensitivity characterization across the line dimension showed a slightly higher phase noise floor at the edges of the line illumination due to the lower intensity and SNR. A Powell lens ^48^ may be considered instead to generate a line profile with uniform intensity distribution. Lastly, though the *en face* LSO image of photoreceptors was very helpful in optimizing the focus of OCT, the same illumination was used for both OCT and LSO, preventing their simultaneous operation due to the intense reflection from the OCT reference arm saturating the LSO sensor. In the future, it will be straightforward to incorporate different wavelengths for both to allow simultaneous LSO and OCT operation.

In conclusion, we demonstrate the first of its kind high-speed line-scan spectral domain OCT equipped with AO that allows high-resolution volumetric imaging and phase-resolved acquisition of light-induced functional changes from cone photoreceptors. The key features – high speed, anamorphic detection, AO or non-AO operation – summarized above are fundamental to the application of this novel technology for interferometric imaging of neural activity in photoreceptors and holds promise for future applications to other retinal cells in general. While this work represents an important step forward in the technology for all-optical, non-invasive interrogation of retinal activity, understanding the underlying mechanisms^15,39,45^ of such optoretinograms is key for its application as a biomarker to study basic visual processes and retinal health in disease and therapies.

## Materials and Methods

### Line-scan retinal imager with adaptive optics (AO)

The detailed system layout is shown in Figure 7a. The top inset shows the measured full width half maximum (FWHM) of the axial PSF of the system, equal to 7.6 μm at an axial depth of 480 μm. This is similar in magnitude to the FWHM of the coherence function computed theoretically (7.7 μm) for the SLD source spectrum (λ_o_ = 840 nm, Δλ = 50 nm, double-humped Gaussian spectrum). A reflective collimator (RC) was used to provide an effective beam diameter of 4 mm for the OCT channel. A plate beamsplitter divided ~30% and 70% of the OCT illumination into the sample and reference arms respectively. A cylindrical lens CL1 (effective focal length = 100 mm) was placed to generate a line field of 4 mm at the entrance pupil (P4) of the system. In Figure 7a, P4 corresponds to the pupil plane Pcirc in Figure 1. An adjustable pupil iris was placed at the entrance pupil to control the beam size. A 1-D galvo scanner (GS), a deformable mirror (DM) and the eye’s pupil (P1) were optically conjugated using achromatic lens-based afocal telescopes - L1 (L_A_ in Figure 1)/L2, L3/L4, L5/L6. The beam size at the eye’s pupil was 4 mm and generated a 2 deg line field on the retina. The power was 3.2 mW at the eye’s pupil, which is well below the maximum permissible exposure for extended retinal illumination.

**Figure 7:**
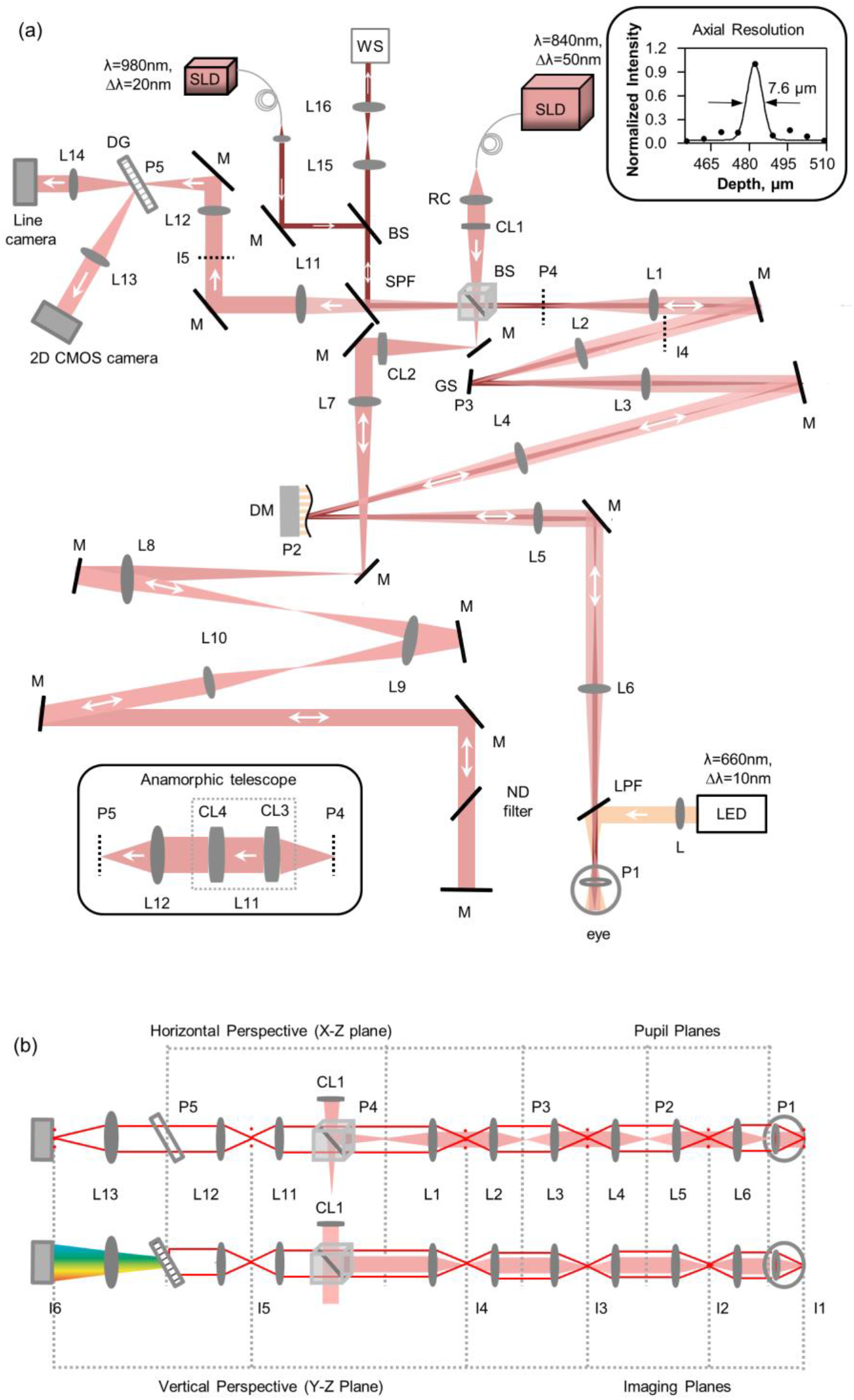
(a) Optical layout of the multimodal line-scan retinal camera, consisting of a line-scan spectral domain OCT and a line-scan ophthalmoscope (LSO) and a visual stimulation channel. The top inset shows the measured axial PSF, the FWHM and best fit. The horizontal perspective (top view) beam path is shown in (a) and the different channels (OCT, Wavefront sensing, and visual stimulation) are denoted with different colors. SLD: superluminescent diode, RC: reflective collimator, CL: Achromatic cylindrical doublet lenses, BS: beam splitter, L: achromatic doublet lenses, M: mirror, GS: galvo scanner, DG: diffraction grating, ND: neutral density, DM: deformable mirror, LPF: long pass filter, SPF: short pass filter, LED: Light emitting diode, WS: wavefront sensor. P: pupil plane, I: imaging plane. The inset shows the anamorphic telescope that replaces lens L11 and described in detail in Figure 1. Focal length of the lenses used are L1=L2=200 mm, L3=L6=250 mm, L4=L5=500 mm, L7=L8=400 mm, L9=300 mm, L10=200 mm, L12=150 mm, L3=200 mm, L14=200 mm, L15=175 mm, L16=125 mm, CL1=CL2=CL4=100 mm, CL3=250 mm, (b) The horizontal (top view) and vertical (side view) perspectives of the optical path are shown. The illumination and detection paths are shown with filled red and solid red respectively. The detection path is shown for the on-axis field point. Note the different beam profiles at the pupil planes in detection compared to illumination.

The reference arm consisted of a second cylindrical lens CL2 (effective focal length = 100 mm) that re-collimated the OCT beam. The collimated beam was then propagated to the final reference arm mirror via three afocal telescopes. L8 and L9 are single achromatic lenses, but the beam propagated through them twice after reflection from the mirrors. The lenses in the reference arm were used for two purposes: first for compensating the dispersion generated in the sample arm due to the afocal lens-based telescopes, and second for reducing the diffraction of the beam generated by long-distance propagation (~ 4 m total double pass from beamsplitter to eye and back). The final mirror of the reference arm was placed in a manual translation stage to match the path length of sample and reference arms to create an interferogram. Note that the beams shown in Figure 7a represent the horizontal perspective (top view), and the detection path assumes a mirror placed at the retina. Below in Figure 7b, we illustrate how the beam profiles differ between the horizontal (top view) and vertical (side view) perspectives, and with illumination and detection.

The backscattered light from the retina propagated via the same path as the illumination. The sample and reference arm beams were combined in the detection arm to generate interference. The beams were propagated via an afocal telescope L11 & L12 (L_B_ in Figure 1), a 1200 lp/mm transmissive grating and a focusing lens L13. An adjustable slit was placed at the image plane I5 (I_anamorphic_ in Figure 1) to provide partial confocality. A fast 2D CMOS camera (Photron, FASTCAM Mini Ax200, pixel size = 20 μm) placed at the focus of lens L13 captured the interferogram. The grating was optically conjugated to the ocular pupil plane. The camera was set to record a region-of-interest of 768 (spectrum dimension, λ) ×512 (line dimension, *x*) pixels. The maximum speed of the camera can be set to 16.2 kHz at this region-of-interest, defining the maximum B-scan or M-scan rate achievable. The lens L11 was replaced with an anamorphic telescope (Figure 6a, bottom left inset) consisting of two cylindrical lenses in order to optimize spatial and spectral resolution simultaneously (Details in Figure 1).

Figure 7b shows the beam propagation cross-sections across the different pupil and imaging planes, in illumination and detection. Note that due to the propagation of the cylindrical wavefront through the system, the beam profile is linear at all the pupil and imaging planes in illumination, but with their orientation rotated by 90 degrees. The line illuminating the retina effectively creates a set of beamlets emanating near-spherical wavefronts when backscattered. Therefore, in detection, the series of linear beamlets create a spherical and linear beam profile at each pupil and imaging plane respectively. The resulting detection path for the on-axis field point is shown in Figure 7b. The backscattered beam overfilled the eye’s pupil leading to a larger effective beam diameter at the pupil planes in detection.

#### Wavefront sensing and correction

A custom-made Shack-Hartmann wavefront sensor and a DM (DM97-15, Alpao, France) were used for measuring and correcting ocular aberrations, respectively. The lenslet array with a pitch of 150 μm and a focal length of 6.7 mm. The Shack Hartmann spot pattern was captured in a CCD camera with pixel size of 6.45 μm. A short pass filter (SPF) with cut-on at 903 nm was used to combine the wavefront sensing (980±20 nm) and OCT beams (840±25 nm). The wavefront sensing illumination followed the same optical path as the OCT sample arm. A smaller beam diameter of 1.5 mm at the eye’s pupil was used and spatially offset from the corneal apex to avoid spurious reflections. In detection, an afocal telescope L15-L16 optically conjugated the entrance pupil (P4), and demagnified the beam to 4.8 mm diameter at the lenslet array. An iris was placed at the focus of L15 to further avoid spurious lens and corneal apex backreflections. A custom-made software allowed closed-loop operation of the wavefront sensor and deformable mirror.

#### Line-scan ophthalmoscope

The LSO used the same path as the sample and the detection arm till the grating. The zeroth-order beam of the grating, which is typically discarded in spectral-domain OCT systems, was used to build an LSO by placing a line-scan camera (Basler, sprint, spL2048-70km, pixel size: 10 μm) with a focusing lens L16. For LSO operation, the intense backreflection from the reference arm was manually blocked. The LSO and OCT cameras were optically conjugated using a model eye. The LSO en face images allowed precise adjustment of the image focus.

#### Visual stimulation

The visible stimulus channel was combined with the remaining beams using a long-pass filter (LPF) (cutoff: 685nm) and introduced into the eye after the last telescope (L5-L6). A 660±10 nm LED was focused at the eye’s pupil with an achromatic lens and provided a homogeneous visual field area of ~37.5 deg^2^. A custom LED driver board was used to control the intensity of the LEDs and to interface timing with the OCT system. The driver board implemented a Pulse Frequency Modulation (PFM) control paradigm and LEDs were mounted on aluminum heat sinks, which taken together, ensured spectral output of the LEDs did not change as a function of intensity. An FPGA running custom VHDL code using an internal clock of 133 MHz generated the PFM and monitored a synchronized TTL pulse generated from the main OCT data acquisition board for when to trigger the stimulus. A maximum delay between OCT TTL signal state change to LED output change was two clock pulses (~15 ns). The pulse width in the PFM paradigm is 2 μs, which exceeds the Nyquist criterion for the shortest pulse duration used for the experiments (5 ms).

##### Acquisition & Processing

For OCT, the 2D images from the camera were stored on the on-board flash memory and transferred via a dedicated ethernet connection to the computer. For LSO, the image stream from the line-scan camera was transferred via a frame-grabber (NI-PCIe-1427)) to the same computer. Synchronization of the galvo scanner, OCT, LSO acquisition, and the visual stimulus was achieved by the analog output board (NI-PCIe-6738). A custom-built software was developed in LABVIEW to display the LSO image stream and real time OCT B-scan reconstruction.

Typical OCT reconstruction steps were followed. Each acquired B-scan was background subtracted to improve the SNR and the phase noise. The background was calculated as the average of 500 images recorded only with reference beam. The background subtracted B-scans in each OCT volume (*I*(*x*, *y*, *λ*)) was resampled (*I*(*x*, *y*, *k*)) and Fourier transformed along the spectrum dimension for each pixel *x* on the line to extract the depth information from the complex numbers *I*(*x*, *y*, *z*).

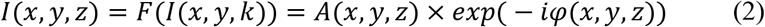

For imaging structure, only the absolute value of the complex number was used. Each OCT volume was segmented using an open-source software ^49^ to obtain *en face* images by taking maximum intensity projections (MIPs) of 3 pixels centered at the cone inner-outer segment junction (ISOS) and cone outer segment tips (COST). The segmentation step also provided axial shifts between different volumes in a time series due to eye motion. The series of *en face* intensity images at the COST layer were spatially registered using a strip-based registration algorithm ^50^ and averaged to improve signal-to-noise ratio (SNR).

For light-induced optical path length changes, each OCT volume was registered using the axial and lateral shifts from the prior step to obtain *I*_*reg*_(*x*, *y*, *z*), and these volumes were used for further analysis. First, each volume was referenced to the mean of all the volumes that were recorded before the start of stimulus to cancel the arbitrary phase.

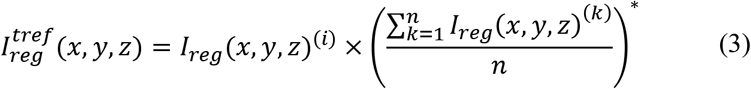

where 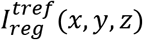 is time referenced complex volumes, i = 1,2..m is volume number (m is the total number of volumes recorded in each measurement), k is the volume recorded before the stimulus, n is the total number of volumes recorded before the stimulus, and * represents complex conjugate. Once referenced over time, the mean of 3 axial pixel complex values, centered at the boundaries of the ISOS, 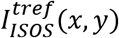 and the cone outer segment tips (COST), 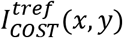 was calculated. The phase difference between these two layers was obtained by first multiplying the complex conjugate of COST with the ISOS layer.

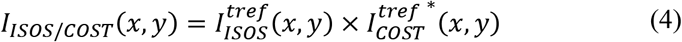

In each volume, the complex numbers *I*_*ISOS*/*COST*_(*x*, *y*) were either averaged over a small retinal area (0.27 deg^2^) for measurements without AO, or averaged over the collection aperture of a cone for measurements with AO. Each individual measurement consisting of one volume series provided ‘m’ complex numbers, converted into a time series, (t), based on the volume acquisition rate ranging between 15 – 324 Hz. For multiple measurements, the complex average of individual time series was obtained. The phase responses (*Δϕ*_*ISOS*/*COST*_(*t*)) were computed by calculating the argument of the complex time series. For phase responses that exceeded ±*π* radians, phase was unwrapped along the time dimension. The change in OPL (*ΔOPL*(*t*)) was calculated as

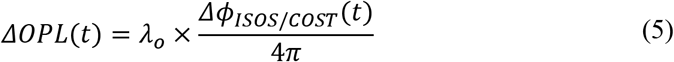

where *λ*_*o*_ is the central wavelength.

#### In vivo imaging protocol

Two cyclopleged (0.5% tropicamide) subjects free of retinal disease participated in the study. The research was approved by the University of Washington institutional review board and all subjects signed an informed consent before their participation and after the nature and possible consequences of the study was explained. All procedures involving human subjects were in accordance with the tenets of the Declaration of Helsinki. Structural and functional imaging of the retina between 2 – 7 deg temporal eccentricity was conducted. A pupil camera was used to monitor the alignment of the ocular pupil. For the light-induced OPL response experiments, the subject was dark adapted for 3 to 4 minutes, and OCT volumes were acquired for 1-3 seconds at different volume rates. For imaging without AO, defocus was optimized either by translating the eye’s pupil and L6 together as a unit, or using the DM, to achieve best focus of cone photoreceptors in the real-time LSO image stream. After 10-20 volumes, the stimulus flash illuminated the retina. The above procedure was repeated under conditions where higher order aberrations were corrected with AO. For the AO ORG experiments aimed at cone spectral classification, 40 volumes were recorded at the speed of 6 kHz B-scan rate and three measurements were averaged.

## Acknowledgements & Disclosures

We thank Daniel Palanker, Austin Roorda, Hyle Park and Yuhua Zhang for helpful discussions. Funding for the study was obtained from NIH grants U01EY025501, EY027941, EY029710, P30EY001730, Research to Prevent Blindness Career Development Award, Foundation Fighting Blindness, Murdock Charitable Trust, Burroughs Welcome Fund Careers at the Scientific Interfaces, Unrestricted grant from the Research to Prevent Blindness.

VPP and RS have a commercial interest in a US patent describing the technology for the line-scan OCT for optoretinography.

